# Deep-learning prediction of gene expression from personal genomes

**DOI:** 10.1101/2024.07.27.605449

**Authors:** Shiron Drusinsky, Sean Whalen, Katherine S. Pollard

**Affiliations:** Gladstone Institutes; San Francisco, CA 94158, USA; Biomedical Informatics Graduate Program, University of California, San Francisco; San Francisco, CA 94158, USA; Chan Zuckerberg Biohub San Francisco; San Francisco, CA 94158, USA; Department of Epidemiology and Biostatistics, University of California, San Francisco; San Francisco, CA 94158, USA

## Abstract

Models that predict RNA levels from DNA sequences show tremendous promise for decoding tissue-specific gene regulatory mechanisms^1–5^, revealing the genetic architecture of traits^6–10^, and interpreting noncoding genetic variation^10,11^. Existing methods take two different approaches: 1) associating expression with linear combinations of common genetic variants (training across individuals on single genes)^12,13^, or 2) learning genome-wide sequence-to-expression rules with neural networks (training across loci using a reference genome)^11,14,15^. Since limitations of both strategies have been highlighted recently^16–20^, we sought to combine the sequence context provided by deep learning with the information provided by cross-individual training. We utilized fine-tuning to develop Performer, a model with accuracy approaching the cis-heritability of most genes. Performer prioritizes genetic variants across the allele frequency spectrum that disrupt motifs, fall in annotated regulatory elements, and have functional evidence for modulating gene expression. While obstacles remain in personalized expression prediction, our findings establish deep learning as a viable strategy.

## Main

As implemented to date, deep learning and linear models provide complementary approaches. State-of-the-art linear models^6,8,9,12^ use penalized regression to identify common variants that predict expression of a single gene across a sample of genotyped individuals with RNA-seq data from one or more tissues/cell types. This strategy can explain much of expression heritability^6,21^. However, these models 1) cannot make predictions for new genes or variants, 2) largely ignore the contributions of rare variants, 3) identify expression quantitative trait locus (eQTL) variants that frequently do not co-localize with trait-associated variants, and 4) require additional fine-mapping and experiments (e.g., epigenetic data) to pinpoint causal alleles^19,20,22^. In contrast, existing deep learning models use a single reference genome per species to predict genome-wide epigenetic and expression data from many tissues/cell types^7,11,14,15^. Feature attribution methods have shown promise for nominating causal alleles including tissue-specifying and disease variants^2,5,10,11,23^. However, existing deep learning models cannot reliably explain expression variation across individuals or predict eQTL direction^16,18^.

Before concluding that neural networks are unable to model inter-individual expression variation, it is critical to apply cross-individual training and evaluation to cohorts of hundreds of individuals where linear models have been successful. Previous work used far fewer individuals and did not evaluate across them.^7,24^ To address this we developed Performer, a fine-tuning strategy that implements cross-individual training and evaluation of sequence-to-expression neural network models. Briefly, we modified the Enformer architecture^15^ by replacing the output head with one that predicts tissue-specific gene expression as a scalar value rather than a genomic track and implemented fine-tuning with Enformer’s weights as starting values for the parameters in the model trunk and a custom loss function (**Methods**). We hypothesized that Performer would retain Enformer’s knowledge of regulatory grammar while gaining the ability to capture expression cis-heritability.

To explore this, we trained Performer models and compared them to Enformer without fine-tuning and to elastic nets, a representative linear modeling framework that performs well at predicting expression variation^6,16,18^ (**Methods**). We obtained paired whole-genome sequencing (WGS) and bulk RNA-seq data from the GTEx study^12^, focusing initially on “Whole Blood’’ samples because of the large sample size (n = 670). We selected ∼300 genes from Enformer’s train set with a range of cis-heritabilities. For each person and gene we generated a 49-kilobase (kb) personalized genome sequence centered at the TSS, including all single-nucleotide variants (SNVs), and paired it with the same person’s normalized blood gene expression value. For each gene, we trained three elastic net and Performer model replicates, matching the genomic window and how we varied individuals used for training and evaluation (**Methods**), enabling us to assess the mean and range of performance. While training Enformer from scratch can take weeks on multiple GPUs, fine-tuning takes only ∼30 minutes per gene on a single GPU.

We first investigated performance on held-out people and train genes (HOP). Using Performer, pre-trained Enformer, and elastic net models, we made blood gene expression predictions for HOP (**Fig. 1a**). As others have done^6,16,18^, we evaluated model predictions in terms of the variance in expression they explain across people with the coefficient of determination (R^2^) and Pearson’s correlation coefficient (PCC). These evaluations show that single-gene Performer models consistently outperform Enformer (**Fig.1a-c**). In agreement with prior studies^16,18^, Enformer’s predictions across GTEx individuals tend to show relatively limited variation or variation that is negatively correlated with observed values. Our fine-tuning corrected this for most genes across the range of expression heritability (**Fig.1a-c; Supp.** Fig. 1). Strikingly, Performer’s metrics are on par with or slightly better than those of elastic nets for 84% of genes. Learning curves generated by downsampling the training set indicate that this level of performance is achievable with 250 or fewer individuals (**Supp. Fig. S2**). Finally, we trained Performer and elastic net models using paired WGS and dorsolateral prefrontal cortex RNA-seq from the ROSMAP study^25^ (n = 742) and made HOP predictions for GTEx individuals with brain cortex RNA-seq (n = 205), observing good cross-cohort performance despite cortex

**Figure 1:**
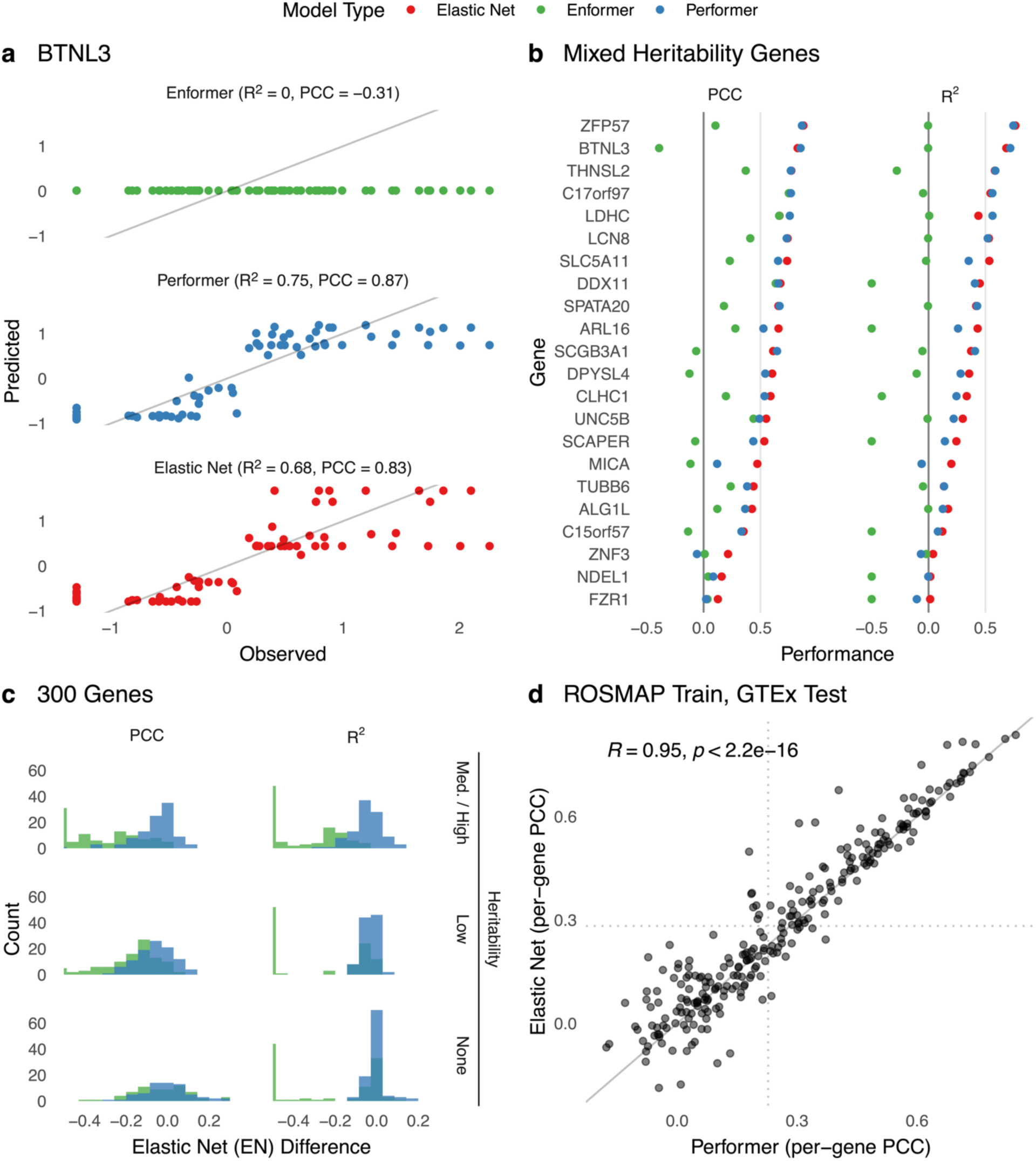
Cross-individual fine-tuning increases expression variation explained by model predictions. All panels display performance on held-out people (HOP) and train genes. a) Coefficient of determination (R^2^) and Pearson correlation coefficient (PCC) between observed expression values and predicted expression using pre-trained Enformer, Enformer after fine-tuning on paired BTNL3 WGS/ Whole Blood RNA-seq data from GTEx (Performer), and a penalized linear model (elastic net) fit on the same data. Each point represents an individual from the test set and their Whole Blood BTNL3 expression. Results are shown for 1 of 3 elastic net and Performer model replicates and the same 71/670 individuals were used to evaluate each model. b) R^2^ and PCC of models trained on single genes with diverse expression cis-heritabilities (as estimated by elastic nets). Performer and elastic net models were trained on single genes and evaluated on the same genes using individuals from the test set, with performance averaged over 3 model replicates evaluated on different splits of individuals. R^2^ and PCC for Enformer are averaged over the same three test sets. Performer and elastic net models are typically within a small margin of each other. Enformer R^2^ values are thresholded at -0.5 for axis visibility. R^2^ is expected to be lower for Enformer (**Methods**). c) Enformer and Performer R^2^ and PCC value for each of 301 genes is subtracted from the elastic net metric evaluated on the identical test set, and the average over 3 replicates is reported. Genes are divided into equal sized groups based on elastic net heritability estimates (R^2^): medium/high (0.06 to 0.76), low (0.01 to 0.06), and none (0 or less). d) PCC of 300 genes using Performer and elastic net models trained on single genes using paired WGS and dorsolateral prefrontal cortex RNA-seq data from ROSMAP and evaluated on the same genes and test individuals using GTEx WGS and Brain - Cortex RNA-seq data (n = 205). Each point represents a different gene. Performance of the Performer and elastic net models is significantly correlated (t-test p<2e-16). PCC values come from one model replicate.

being a less specific brain region (**Fig. 1d**). These results establish that deep neural networks trained on paired WGS and RNA-seq can accurately model the relationship between gene expression and cis regulatory variants. Thus, the gap in performance between prior deep learning models and linear models can be attributed primarily to lack of cross-individual training, not to shortcomings of neural networks per se.

Next, we investigated why Performer achieved competitive performance. We defined High Scoring Variants (HSVs) by ranking SNVs by the absolute difference between each model’s alternate and reference allele expression predictions, then used the number of non-zero elastic net coefficients to establish a consistent HSV set size across models. While elastic net HSVs tend to be distributed over the whole window (**Fig. 2a**), Performer’s and Enformer’s HSVs are generally more TSS proximal (**Fig. 2a**-*ZFP57;* **Fig. 2b**). However, Performer’s HSV-TSS distances vary, with some genes having primarily distal HSVs (**Fig. 2a**-*BTNL3*). This suggests that, for some genes, cross-individual training has helped Performer overcome Enformer’s underutilization of distal variants^17^. Performer and elastic net HSVs largely have the same direction of effect, while Enformer’s scores often disagree (**Fig. 2c**)^16,18^, leading to negative cross-individual correlations (**Supp. Fig. S1**). These findings show that cross-individual training has improved two key limitations of existing deep learning models.

**Figure 2:**
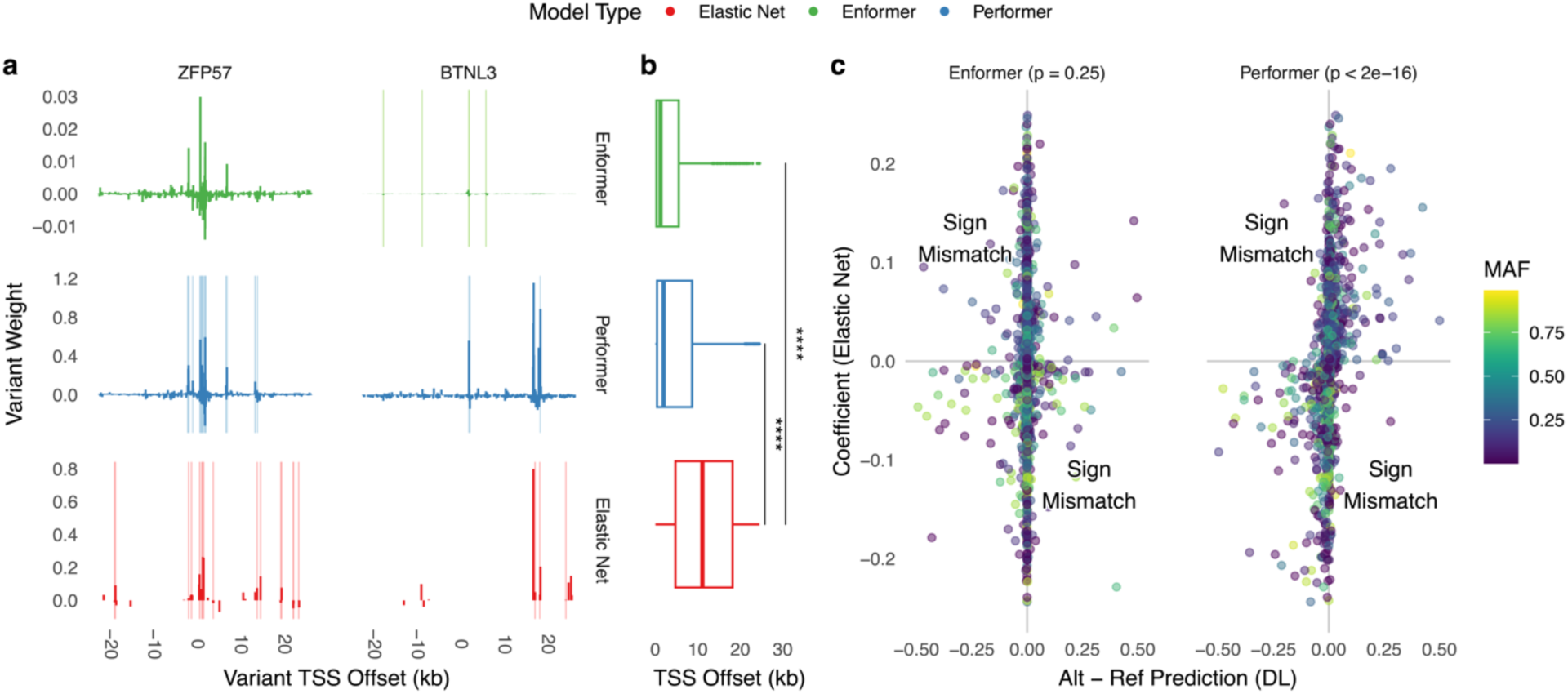
Cross-individual fine-tuning alleviates previously identified limitations of sequence-to-expression models. a) Variant weights (elastic net: coefficient; Enformer/Performer: alternative-reference prediction) versus variant position relative to TSS for *ZFP57* and *BTNL3*. Light vertical lines represent significant motif disruptions at HSVs. b) Distribution of high-scoring variant (HSV) proximity to TSSs for 42/301 genes with Performer R^2^ > 0.2. HSVs for a given gene include all SNVs with non-zero coefficients for elastic net models and the same number of SNVs with the greatest absolute ISM values for Enformer/Performer. c) Variant weights of elastic nets versus Enformer (left) or Performer (right), for the same genes as in (**B**), demonstrating increased support for Performer variant directional effects, regardless of minor allele frequency. *p*-value from Fisher’s exact test. Only SNVs whose elastic net coefficients are non-zero are shown.

We wondered if Performer’s increased similarity to elastic nets would decrease identification of SNVs with functional evidence. Instead, Performer’s HSVs have functional signatures similar to Enformer’s and significantly stronger than elastic nets’, including enrichment in active enhancer and promoter chromHMM states (**Fig. 3a**) and disruption of transcription factor motifs (**Fig. 3b**). Furthermore, Performer’s variant weights predict fine-mapped GTEx eQTLs on par with elastic net’s and better than Enformer’s (**Fig. 3c**). Thus, cross-individual, single-gene training does not notably degrade the neural network’s encoding and weighting of regulatory sequence patterns.

**Figure 3:**
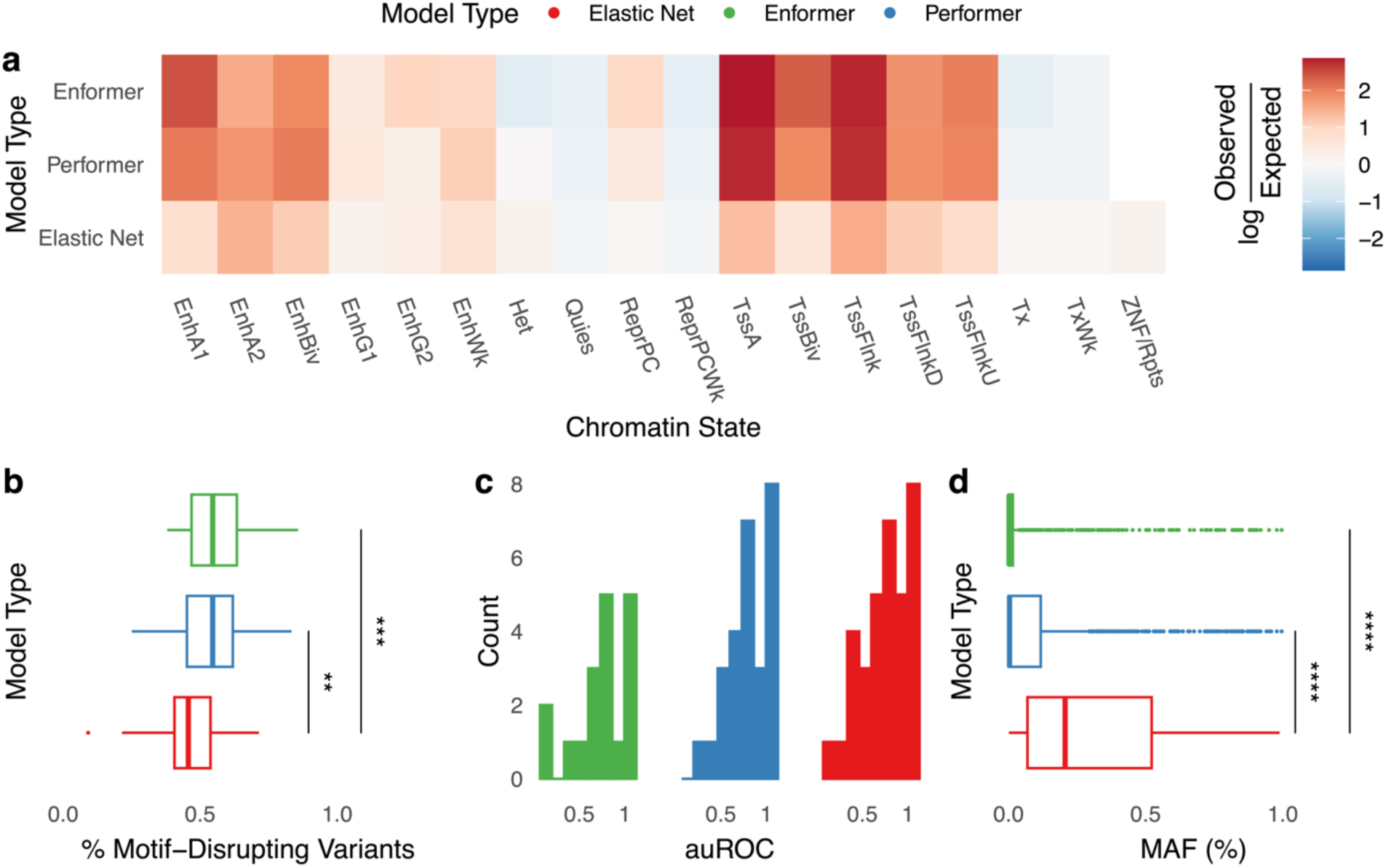
Cross-individual fine-tuning retains weighting of regulatory sequence patterns. All panels display properties of Enformer, Performer, and elastic net HSVs for 42/301 genes with Performer R^2^ > 0.2, evaluated on held-out people (HOP) and train genes. a) Enrichment of HSVs across ChromHMM^36^ chromatin states in monocyte cells from Epimap^37^, averaged across 3 donors. b) The percent HSVs that significantly increase or decrease the affinity of an overlapping DNA-binding protein’s motif, as measured by *motifbreakR*^34^ with a p-value threshold of 5e-5. c) Distribution of area under the Receiver Operating Characteristic (auROC) curve for predicting CAVIAR^38^ fine-mapped Whole Blood GTEx eQTLs. HSVs were labeled as being fine-mapped eQTLs or not, and this label was predicted based on thresholding absolute variant weights from each model (Enformer/Performer: ISM; elastic net: coefficients). The auROC was computed for each gene, and genes were dropped if no HSVs were fine-mapped eQTLs (Elastic Net: 6, Performer: 13, Enformer: 22). Performer and elastic net models have similar distributions of auROC, while Enformer auROC is worse as expected since the model was not trained on inter-individual variation. d) Distribution of HSV minor allele frequencies (MAFs).

Since deep learning models tend to score rare variants higher than common eQTLs^23^ and the minor allele frequencies (MAFs) of Performer/Enformer HSVs are generally lower than those of elastic nets (**Fig. 3d**), we repeated the functional signature analyses after removing HSVs with MAFs <1% or <5% and found that Performer’s functional enrichments persist (**Supp Fig. S3**). For example, the top Performer HSVs of *ZFP57* include both rare variants (**Fig. 4a,c**), and common variants (**Fig. 4b,d**) that create stronger matches for blood transcription factor motifs. Elastic nets upweight adjacent SNVs that do not alter motifs and assign zero weight to this common variant. Next, we identified a small number of driver variants for each Performer/Enformer model by creating a surrogate linear model with comparable performance to the full model via forward selection^16^ (**Fig. 4e**, **Supp. Fig. S4**). Compared to all HSVs, drivers have somewhat higher MAFs and lower pairwise LD (**Supp. Fig. S4**). Nonetheless, they maintain the functional properties of HSVs (**Fig. 4f-g**). These results show that Performer’s weights prioritize SNVs with functional signatures across the allele frequency spectrum, including variants that explain gene expression variability. Thus, Performer has useful features previously associated with both linear and deep learning models.

**Figure 4:**
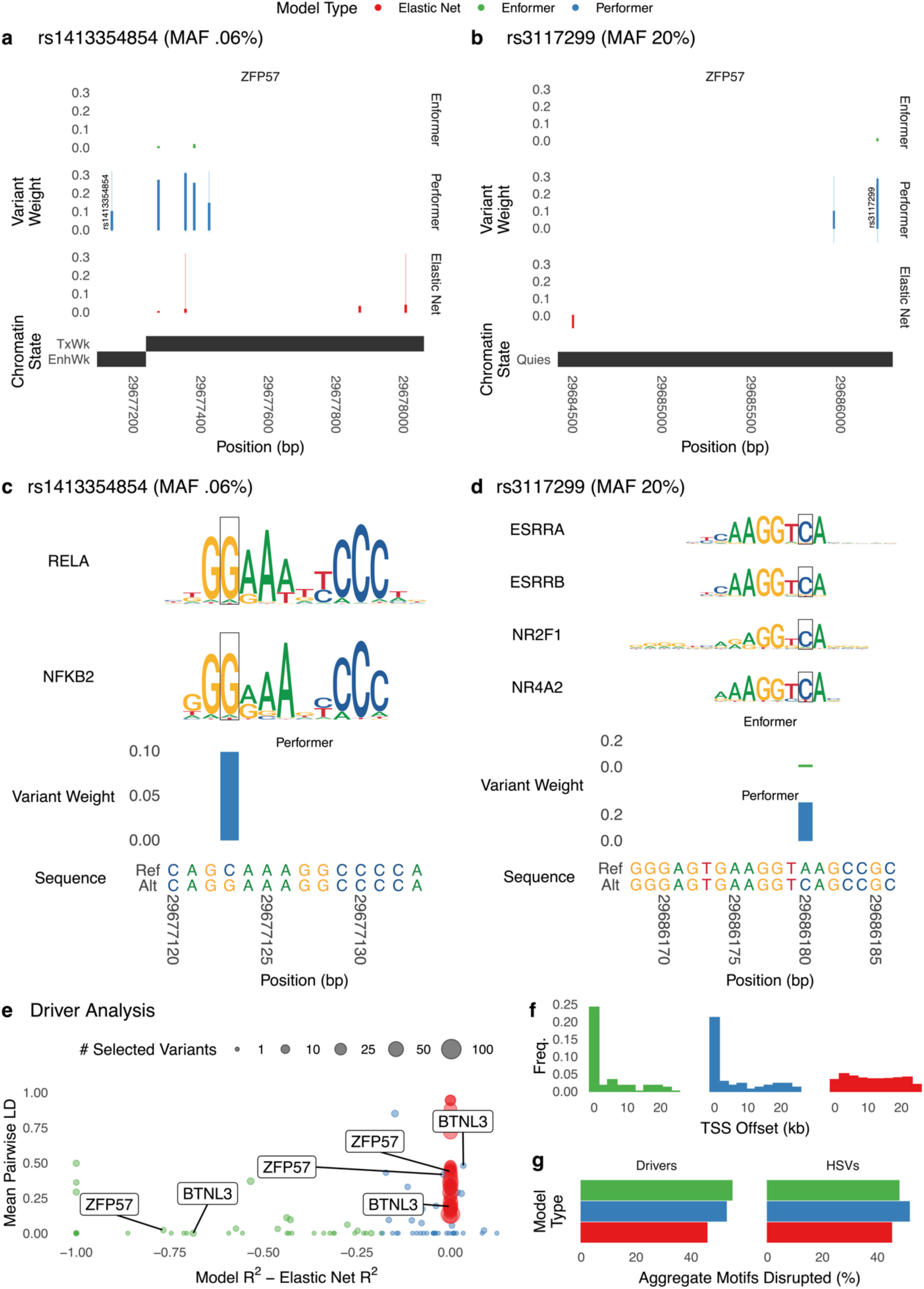
Performer identifies HSVs that alter motifs and drive expression predictions. a) Genomic track view of variant weights (Enformer/Performer: ISM (**Methods**); elastic net: coefficient) around a rare HSV (rs1413354854; GTEx MAF=0.06%) of *ZFP57* that falls in a ChromHMM^36^ weakly active enhancer. Performer highly weights the HSV, while Enformer and elastic net assigns moderate weights to several flanking SNVs. For each model, wide bars denote variant weights, and thin lines denote positions of HSVs. b) Genomic track view of Enformer, Performer, and elastic net variant weights around a common HSV (rs3117299; GTEx MAF=19.98%) of *ZFP57* that falls in a quiescent chromatin region. Performer highly weights the HSV, while elastic net and Enformer have no HSVs nearby. For each model, wide bars denote variant weights, and thin lines denote positions of HSVs. c) Zoomed in view of rs1413354854 showing that the alternative allele creates a binding site for NFKB2 and/or RELA. Logos generated using *motifbreakR*. d) Zoomed in view of rs3117299 showing that the alternative allele creates a binding site for several blood transcription factors. Logos generated using *motifbreakR*. e) Model performance on HOP relative to elastic net plotted against the mean pairwise linkage disequilibrium (LD) of selected variants (Enformer/Performer: drivers (**Methods**), elastic net: non-zero coefficients), with the number of selected variants indicated by dot size. For the vast majority of genes, a very small number of Performer/Enformer driver variants are identified (resulting in a lower mean LD) compared to penalized regression with elastic nets, though forward selection could similarly be used with a non-penalized model to select a smaller number of predictive variants. Only 42/301 genes with Performer R^2^ > 0.2 are shown. f) Distribution of selected variant distance from TSS (Enformer/Performer: drivers, elastic net: non-zero coefficients). g) Rates of motif affinity changes based on *motifbreakR* for HSVs and selected variants (Enformer/Performer: drivers, elastic net: non-zero coefficients).

Given these characteristics, we hypothesized that training Performer models on multiple genes simultaneously might improve their performance. A multi-gene model based on all ∼300 training genes outperformed Enformer on most of these genes (**Fig. 5a**) but was comparable to single-gene models (**Fig. 5b**), when evaluated on HOP. We wondered if multi-gene models might be better at generalizing to genes not used in training Enformer or Performer, reasoning that information integrated from various loci would offer the best chance at success for this task. Enformer and multi-gene Performer explained a similar amount of inter-individual gene expression variability on held-out genes and people (HOGP) (**Fig. 5c**), which is much less than single-gene models trained directly on the same genes (**Supp. Fig. S5**). Increasing multi-gene Performer’s training set to ∼11,000 genes did not boost performance on HOP or HOGP (**Supp. Fig. S5**), nor did 196-kb sequences (**Supp. Fig. S7**). Thus, the level of generalizability required to predict causal variant effects in novel loci remains an important open challenge, as this is expected of models that have comprehensively learned causal regulatory patterns.

**Figure 5:**
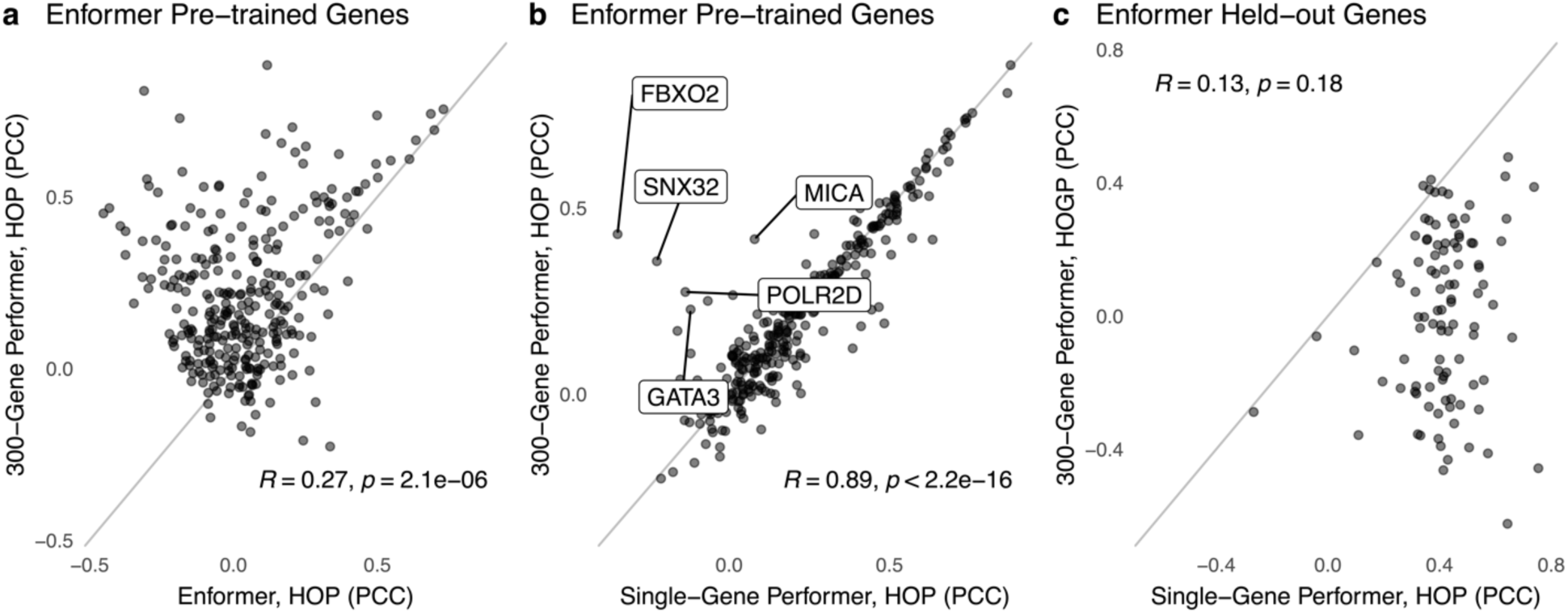
Multi-gene training does not increase expression variation explained. Each point represents the performance of a model on a single gene using GTEx Whole Blood expression values. Results are not averaged over replicates as in (Fig. 1B); they come from one model and one evaluation set only. a) PCC of Performer jointly trained on 301 genes (y-axis) versus Enformer (x-axis), both evaluated on each of the 301 training genes. For most genes, Performer explains more expression variability. Evaluations were performed on held-out people (HOP) and train genes. b) PCC of Performer jointly trained on 301 genes (y-axis) versus Performer trained on single genes (x-axis), both evaluated on each of the 301 training genes. Performance is very similar for multi-gene and single-gene models. A few exceptions, for which multi-gene training improved performance, are labeled. Evaluations were performed on HOP and train genes. c) PCC of Performer jointly trained on 301 genes and evaluated on 100 held-out genes and people (HOGP; y-axis) versus Performer directly trained on each of these 100 genes and evaluated on held-out people (HOP; x-axis). The 100 genes used in both evaluations have high expression cis-heritability and were not used to train Enformer or Performer (**Methods**). For most genes, multi-gene fine-tuning did not achieve the same PCC on unseen genes as can be achieved by directly training on the gene.

We next investigated whether making different modeling decisions would change our findings. We expanded our fine-tuning strategy to the Borzoi model^26^, which has a different architecture from Enformer, was pre-trained directly on RNA-seq rather than CAGE-seq, and accepts longer sequences (524 vs 192 kb) while outputting higher resolution tracks (32 vs 128 bp). Borzoi models fine-tuned on GTEx blood RNA-seq (scalar values) achieve comparable HOP performance to Performer (**Supp. Fig. S6**). We also fine-tuned Enformer and Borzoi on blood RNA-seq read coverage tracks. Both models capture observed coverage variability at gene promoters and ignore it at intergenic regions, but they sometimes under-estimate inter-individual expression differences, albeit not at the same locations (**Fig. 6**). Thus, both architectures perform well at de-noising but may mistake technical and biological variation under different circumstances. This suggests that further minor improvements to sequence length, output resolution, expression data encoding, and architecture may not offer significant performance gains over linear models for cross-individual expression prediction.

**Figure 6:**
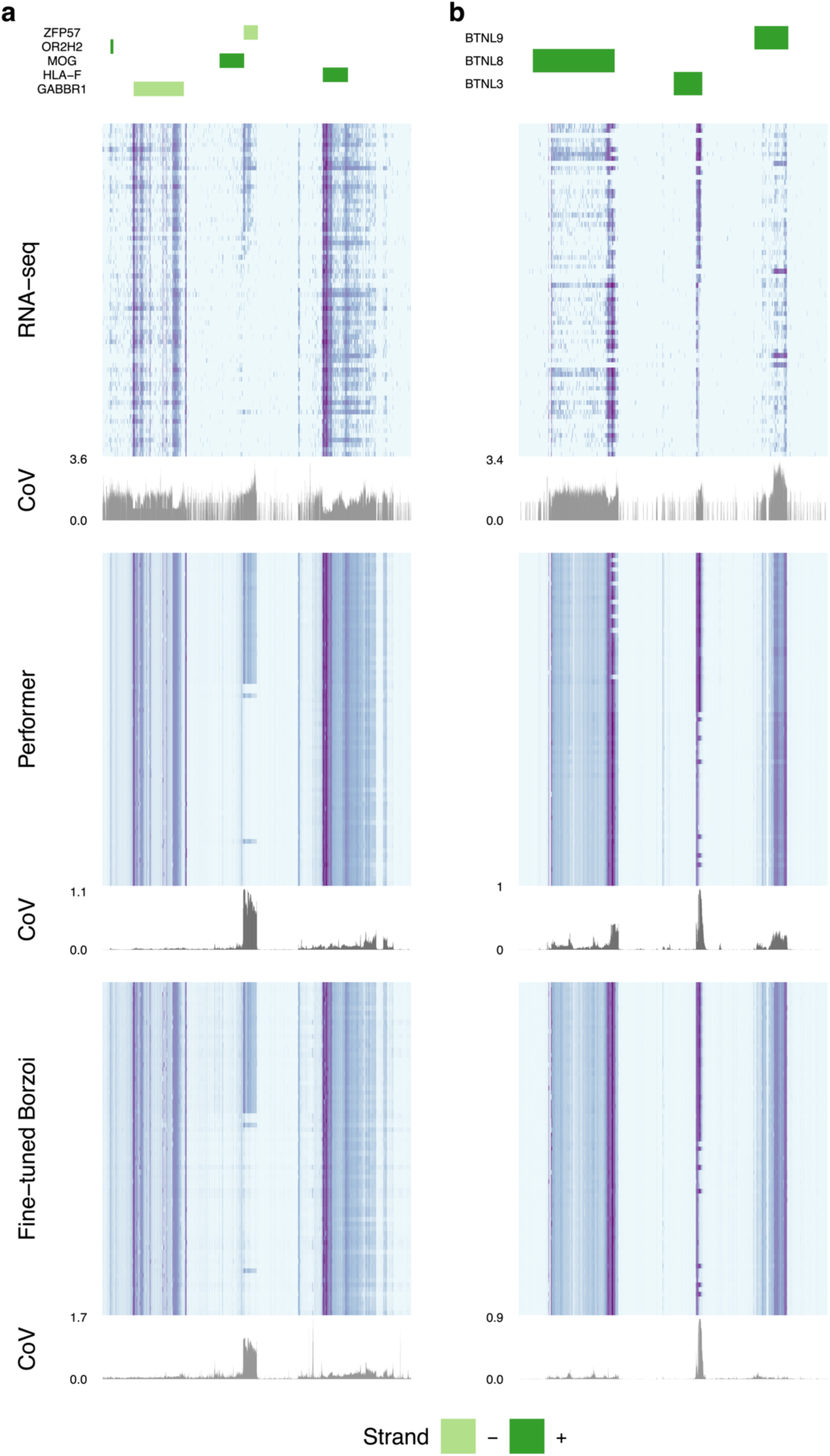
Fine-tuning with alternative architectures. a) Genomic region centered on the *ZFP57* gene showing flanking genes (top), observed RNA-seq coverage for HOP (second from top; purple: high coverage, light blue: low coverage) with its coefficient of variation (CoV), predicted coverage on the same HOP using fine-tuned Enformer (Performer; middle heat map, same color scale) and its CoV, and predicted coverage using fine-tuned Borzoi (U-net architecture; bottom heat map, same color scale) and its CoV. Rows of heat maps correspond to different held-out individuals, sorted in all three heat maps from highest to lowest expression at the *ZFP57* TSS. Both models predict de-noised versions of the observed RNA-seq signal while generally capturing predictive, high CoV regions across people. Fine-tuned Borzoi captures a high CoV region downstream from the ZFP57 TSS that Performer misses. b) Genome region centered on the *BTNL3* gene showing the same analyses as in (a). Performer captures expression variance at *BTNL8* and *BTNL9* that is missed by fine-tuned Borzoi. Both models capture high CoV regions at *BTNL3* and a region containing the HSV described in (Fig. 2A).

In conclusion, we showed that training on personalized genome data and paired expression from hundreds of individuals dramatically improves deep learning models’ ability to predict expression differences amongst unseen individuals. This clarifies that cross-individual training is key to personalized gene expression imputation, and that neural networks can identify SNVs that drive competitive performance at this task. In terms of causal variant prediction, Performer retains Enformer’s utilization of SNVs in candidate regulatory elements, while gaining linear models’ ability to correctly predict the direction of SNV effects on gene expression and to prioritize fine-mapped eQTLs. Additionally, Performer HSVs include both rare and common variants. These characteristics suggest that Performer variant weights hold promise for causal variant detection, though future high-throughput experimental validations are needed to test this hypothesis.

The Performer approach has several important limitations. First, similar to eQTL models, Performer only estimates the genetic component of gene expression variability and performs best on genes with high cis-heritabililty. Second, we only investigated SNVs in this study; it will be important to extend Performer to insertions, deletions, and inversions, because they are major contributors to expression variability^27^. Third, Performer is competitive with existing models at complementary tasks (e.g., using common variants to impute expression of unseen individuals for elastic nets, functional variant prioritization across the MAF spectrum for Enformer), bridging the gap between these modeling approaches. However, it does not out-perform either method and hence has room for improvement. Furthermore, fine-tuning did not address the ambitious task of predicting the expression variability of unseen genes (which no model currently does), even when trained jointly with thousands of genes or larger input sequences. This lack of generalizability may be due to Performer overfitting to specific eQTL associations in training genes, as evidenced by decreased ability to predict average expression of unseen genes (**Supp. Fig. S8**), or upweighting changes in motifs that have limited or even opposing effects on other genes’ expression.

Our results suggest several future directions that may help sequence-to-expression models realize their full potential to decode gene regulatory logic. For example, training efforts may need to account for the strongest eQTL signals, such as by masking them during training, to help models better attend to the remaining patterns and avoid memorization of eQTL positioning. Performance may also be bolstered by combining cohorts like GTEx with massively parallel reporter assay (MPRAs) and high-throughput genome editing perturbation data, which includes many rare and de novo variants. Indeed, others have suggested the variation present in the human genome is insufficient for learning causal patterns from sequence^28^; larger synthetic MPRA libraries may help bridge this gap. New architectures and loss functions designed to capture biological variability are also worth exploring. Nevertheless, incorporating genetic variation via fine-tuning corrects previously documented issues for cross-individual expression prediction and demonstrates the promise of deep learning for the field.

## Methods

### Inputs

For each individual in ROSMAP and GTEx, we used *bcftools consensus*^29^ to replace reference genome (hg38) nucleotides with single-nucleotide variants (SNVs) present in their unphased VCF file. All SNVs were used, without allele frequency filtering, to standardize the genetic variation seen by each approach. This led to two consensus sequences per individual; one contains alternate alleles only for homozygous SNVs and reference alleles elsewhere, the other contains alternate alleles for heterozygous SNVs and homozygous SNVs. If phased haplotypes were available, Performer could work with those.

During training and evaluation, we used 49,152 basepair (bp) input sequences that are 4x shorter than Enformer is equipped to handle (196,608 bp) to enable faster and larger batches within the available GPU memory. For each gene, we fetched an individual’s two 49-kilobase (kb) consensus sequences centered on the gene’s TSS (GENCODE^30^ v26). We one-hot encoded each sequence, and used the average of two one-hot encoded matrices as our input^24^. While other approaches are possible^7^, we found this representation to be reasonable because:

(1) it allows for one input per person, speeding up training and evaluation; (2) since the variants are unphased and not matched to allele-specific expression, it avoids the situation where the model is trained or evaluated on two different sequences from the same person being paired with the same gene expression target; (3) the resulting encoding represents whether a person has no dosage (0/0; 0), half dosage (0/1; 0.5), or full dosage (1/1; 1) of any particular SNV.

### Targets

For GTEx “Whole Blood” and “Brain - Cortex” expression, we downloaded tissue normalized data from GTEx^12^ v8. ROSMAP TPM values were normalized using the same procedure (https://gtexportal.org/home/methods; https://github.com/broadinstitute/gtex-pipeline/) During training and evaluation, each individual’s tissue-specific gene expression value is represented as a scalar target value.

### Fine-tuned Enformer Architecture and pre-trained weights

We fine-tuned the pre-trained pytorch implementation of Enformer (https://github.com/lucidrains/enformer-pytorch), using the provided HeadAdapterWrapper class, with num_tracks set set to 1, post_transformer_embed set to False, and the output activation set to the identity function. This bypasses the original human and mouse Enformer heads, passing instead through a linear layer with no activation. The output after passing in a 49,152bp DNA sequence is length 384. We converted this sequence into a scalar by keeping one of the TSS-overlapping center bins (bin 192). This value was used to compute the loss during training, and was compared against observed expression values during evaluation.

### Definition of TSS-centered genomic intervals for training and evaluation

We retrieved the genomic intervals corresponding to training, tuning, and testing for Enformer^15^. First, we downloaded the Basenji2^31^ genomic intervals from https://console.cloud.google.com/storage/browser/basenji_barnyard/data. Next, we extended these 131,072 bp intervals to 196,608 bp. Although this was not done in Avsec et al., we additionally dropped genomic intervals from the train set if, after extending the sequence length, any of these intervals leaked into the tuning set or test set. Likewise, we dropped intervals from the tuning set if extending them caused leakage into the test set.

Of these remaining genomic intervals, we next found 196,608-bp regions that are centered on the TSS of a protein-coding gene and are completely contained within either the train, tune, or test set. We used GENCODE^30^ v26 annotations to define TSS coordinates of protein-coding genes. We omitted from consideration 228 train, 21 tune, and 36 test genes that were within +/-64 bp of another gene’s annotated TSS. We made this decision because DNA sequences from these genes’ genomic intervals would include the TSS for multiple genes within the same Enformer output bin (each bin along the sequence axis of Enformer’s output represents 128 bp of original sequence). We kept only genes with sufficient expression (see **Targets**), leaving us with 11,429 train, 1057 tune, and 1398 test genes for models to be trained on GTEx Whole blood and 12,292 train, 1194, tune and 1492 test genes for models to be trained on ROSMAP DLPFC. When training or evaluating 49,152-bp sequences, we trimmed the left and right edges of these 196,608bp regions to keep only the center portion of the sequences containing the annotated TSS.

### Selection of train and test genes

To select train genes, we first fit elastic nets (separately for ROSMAP DLPFC and GTEx Whole Blood) on approximately 1000 genes in Enformer’s original train set, and computed R^2^ using one cross-validation fold (see **Training Performer Models** and **Fitting Penalized Linear Models**). We then split the resulting R^2^ distribution into 10 equally sized bins and attempted to select 30 genes from each bin, starting with the largest R^2^ bin. If there were less than 30 genes available, more would be sampled from the next bin to compensate. For each of ROSMAP and GTEx Whole Blood, this left us with 300 genes whose elastic net R^2^ approximate the underlying distribution of genes in that tissue, while keeping as many high heritability genes as possible given training time. We added an additional gene, *PPIF*, to the whole blood train set, because of the availability of published perturbation data around this locus measured in THP-1 cells^32^, to enable future experiments that benchmark variant weights from different models against published perturbation data at this locus. In total, this generated 301 genes for training models using GTEx Whole Blood data and 300 genes for training models using ROSMAP DLPFC data.

To select a subset of test genes, we similarly fit elastic nets on every gene in Enformer’s validation (i.e., tuning) and test set, and selected up to 100 genes whose elastic net R^2^ was at least 10%. This led to 82 test genes in ROSMAP DLPFC and 100 in GTEx Whole Blood. We prioritized test genes with strong elastic net performance so we would be able to confidently detect accurate predictions (measured by PCC or R^2^) if they occurred. These are the held-out genes in Figures 5 and S5.

For our experiment involving training on 11,429 train genes, we selected these genes by fetching all genes in the train set (see **Definition of TSS-centered genomic intervals for training and evaluation**).

### Training Performer Models

We fine-tuned Enformer with a learning rate of 5e-6, bf16-mixed precision, and clipping the gradient’s global norm to less than 0.05 using Pytorch Lightning (https://lightning.ai/docs/pytorch/stable). We fine-tuned with 49,152-bp sequences and a batch size of 32 on an H100 GPU. We used gradient accumulation for 4 batches to achieve an effective batch size of 128. All samples in each batch corresponded to data from different people but the same gene, and the same gene was used for all batches until the 4 iterations of gradient accumulation had completed. We chose to include only different people but the same gene in each batch because our loss function emphasized small expression differences between people (see **Loss Function**) to help learn the subtle impact of genetic variants on expression.

An epoch was defined as a step through all genes in the train set and the maximum number of unique individuals that fit into an integer multiple of the effective batch size of 128, which was 512 individuals for both the GTEx Whole Blood and ROSMAP training sets. For single gene Performer models, an epoch would end once these individuals were stepped through, and the tuning epoch would then proceed.

We performed 3-fold cross validation by splitting the total number of individuals with matched WGS and RNA-seq data into train/tune/test splits. For models trained on GTEx Whole Blood data, this was an 80/10/10% split of 670 individuals. Models trained on ROSMAP DLPFC data were trained using a 90/10% train/tune split of 742 individuals and then tested on 205 unseen individuals for which there was WGS and GTEx Brain - Cortex RNA-seq data.

We evaluated the average R^2^ on all genes in the train set and individuals from the tuning set every epoch, and implemented early stopping if this value did not increase with a patience of 20 epochs. After training, the model checkpoint with the greatest average R^2^ on the tuning set, among genes in the train set, was used for evaluation with the test set and subsequent experiments. The test set included test set individuals (described above) and test set and/or train set genes, depending on the experimental goal.

To train jointly on 11,429 genes, we used the “ddp_find_unused_parameters_true” strategy in Pytorch Lightning to train with 8 H100 GPUs. To support our loss function, which emphasizes differences between people for a given gene (see below), we implemented a custom sampler to divide these genes equally among the 8 GPUs and ensure that a batch received by any device always includes data from different individuals but for the same gene. We used a batch size of 16, and evaluated the model on all train genes and individuals from the tuning set every 2 epochs. We stopped training after 33 epochs (7 days), when PCC evaluated on train genes and tuning individuals stopped increasing. We trained only one model due to the computational resources required.

Our “tuning” set is equivalent to what is commonly referred to as a “validation” set. We choose to use the word “tuning” to avoid confusion between the meaning of “validation” in the contexts of machine learning and biological experiments. In the former, it is an intermediate dataset used to estimate performance during training and avoid over-fitting. In the latter, it is often referred to as the final experimental group used to confirm results. Here, the tuning set is used to estimate performance during training, while the test set is the name of the final set of held-out samples used to evaluate trained models.

### Loss Function

Our loss function contained two components. First, the mean-squared error between observed and expected prediction values. Second, we took the pairwise difference between each unique pair of observed expression values within the batch, and performed the same operation for predicted expression values. We then took the mean squared difference between these two pairwise difference vectors. Minimizing the first component incentivizes predictions to be similar to observed values. Minimizing the second component incentivizes the predicted difference in expression between different people to be similar to their true differences. Because all samples in a batch corresponded to different people but the same gene, we reasoned this would emphasize differences conferred by genetic variants onto gene expression, as opposed to differences in expression between different genes. Finally, we took the weighted sum of the two components, weighting each component by one half.

### Calculation of Pearson correlation coefficient and R^2^ across people

We applied the same three cross-validation splits (see **Training Performer Models**) to elastic net and Performer models, thereby maintaining consistency between the individuals used to train and evaluate them for different model replicates. For example, Performer and elastic net models based on the first cross-validation split used identical train/tune/test individuals, and likewise for replicates based on the other two splits. We evaluated Enformer on the same test sets. Unless otherwise specified, for all three model types, we report Pearson correlation coefficients (PCCs) and R^2^ (coefficient of determination) obtained from the average of these three test sets for when evaluating on GTEx Whole Blood. When evaluating on GTEx Brain - Cortex, model replicates were not trained to conserve computational resources. Thus, one model per gene is trained on ROSMAP and evaluated on 205 individuals with GTEx Brain - Cortex WGS & RNA-seq data. We report Enformer’s performance on the same 205 individuals. Unless otherwise specified, we used 49kb genomic windows to train and evaluate elastic net models and Performer models, and evaluate Enformer models, to facilitate further consistency. We either evaluate models on held-out people and the genes they were trained on (HOP) or, for Performer and Enformer, we also evaluate on held-out people and genes (HOGP). Unless otherwise specified, analyses were conducted on HOP.

The PCC and R^2^ across people were calculated separately for each gene through the following procedure. For a given gene, we created a vector of observed gene expression values for all individuals in the test set as well as a second vector of predicted values for the same individuals, in the same order. For the fine-tuned Enformer models (Performer), the predicted values were generated by passing in each person’s DNA sequence for that gene, and the scalar output was used (see **Fine-tuned Enformer Architecture and pre-trained weights** for more details on obtaining a scalar output). For Enformer, we compared the ‘CAGE:blood, adult, pool1’ (track 4950) output to observed GTEx Whole Blood expression values, and the ’CAGE:brain, adult’ (track 4980) output to observed GTEx Brain - Cortex expression values, as others have done^16^. We took the sum over the three center bins of the output, and this value was added to the second vector of predicted values. For 49,152-bp input sequences this was bins 191, 192, and 193 out of 384. We note that the R^2^ for Enformer is expected to be lower, as this model was trained to predict count values rather than normalized TPM values; Enformer will be disadvantaged with R^2^ even if it ranks individuals’ expression values correctly, which will be captured by PCC. Finally, for elastic nets, the predicted values were obtained by passing in each test individual’s genotype matrix to a model fit on the same gene, including only SNPs within the 49,152bp region centered on that gene’s TSS. Using these vectors of observed and predicted expression values, we computed correlation using *scipy.stats.pearsonr* and R^2^ using *sklearn.metrics.r2_score*. We note that PCC is not the square root of R^2^ for non-linear models, motivating us to report both performance statistics.

### Driver Analysis and *in silico* Mutagenesis

For any given gene, we performed *in silico* mutagenesis (ISM) using fine-tuned models and base Enformer by first performing one prediction with the TSS-centered reference genome sequence (49,152 bp). Then, for each observed SNV within the input window in any individual in GTEx, we took the same sequence and replaced the reference nucleotide with the alternate allele nucleotide at the same position, and performed another prediction. For Enformer, we kept predictions from the 3 central bin positions (bins 191, 192, and 193), performed a sum over these bins, and the resulting value was treated as the final prediction. For Performer models, because they were optimized to only use the center bin position (bin 192), we kept only this value. For each SNV, the ISM score (variant weight) was defined as the predicted value from the sequence with the alternate allele minus the one from the reference sequence. Absolute scores were used in analyses that rank SNVs, while the signed ones were used for comparison to elastic net coefficients.

To identify driver SNVs, defined as those that linearly approximate the model’s predictions for a given gene in a desired group of people, we followed a similar forward selection procedure to that in Sasse et al.^16^ To identify drivers, we used only individuals from the test set (i.e., the held-out set of ∼70 people for Whole Blood GTEx models or the 205 individuals with WGS and GTEx Brain - Cortex RNA-seq data for ROMSAP models), to identify variants that explain R^2^ and PCC in these individuals. We deviated slightly from the Sasse et al. procedure by including driver SNVs, as opposed to iteratively including all SNVs, to the linear approximation while searching for drivers.

### Fitting Penalized Linear Models

Linear models were fit using a scikit-learn^33^ pipeline consisting of a VarianceThreshold transform followed by an ElasticNetCV or LassoCV model with max_iter increased to 2000. Models were fit on GTEx or ROSMAP BCF files encoded as a genotype matrix with homozygous reference = 0, heterozygous = 1, and homozygous alternate = 2. We fit and evaluated these models using the same cross-validation folds as Performer.

### Selection of HSVs

To improve variant comparisons across model types, the number of non-zero linear model variant coefficients per gene was computed. An equal number of variants per gene was selected for Performer and Enformer using the highest scoring variants defined by the absolute value of the alternate - reference allele prediction. Analyses using these high scoring variants are thus comparing an equal number of variants per gene across model types. Performer and elastic net HSVs come from those trained on the first cross-validation split.

### Borzoi Analysis

Transfer learning was used to train single-gene Borzoi^26^ models to predict scalar normalized expression values across people. Embeddings were fed into a linear output layer trained with learning rate 1e-3 and early stopping. Separately, fine-tuning was used to train single-gene Borzoi models to predict expression tracks (rather than single values) across people. Output layers were replaced with a dense layer having softplus activation. The Borzoi trunk was initially frozen for 5 epochs to update the new output layer weights, then all layers were made trainable for 20 epochs with early stopping.

### MotifbreakR analysis

Variant-induced changes to the binding affinity of TF motifs was calculated using motifbreakR^34^ with HOCOMOCO^35^ v11 and a p-value threshold of 5e-5.

### ChromHMM analysis

Enrichment of HSVs for epigenetic states was computed using ChromHMM^36^ annotations from Epimap^37^, averaged over 3 peripheral blood mononuclear cell samples. HSVs for each model were overlapped with state annotations, creating an observed number of overlaps per gene/state. An expected number of overlaps per gene/state was computed by first dividing the total length of a state’s annotations that overlap the 49kb gene window by the total length of all annotations overlapping the window. This ratio was then multiplied by the number of HSVs per gene. Enrichment per model/state was defined as the log of the observed over the expected number of states.

### auROC analysis

CAVIAR^38^ high-confidence fine-mapped whole blood eQTLs (FM-eQTLs) from GTEx were used to annotate HSVs. For each model type and gene with at least 1 FM-eQTL, we computed the area under the Receiver Operating Characteristic (auROC) curve using the FM-eQTL annotation and the absolute value of the model’s variant weight. For each model, genes with no FM-eQTLs in its HSVs were omitted as auROC is undefined.

### Training with downsampled donors

For each cross-validation fold (see **Training Performer Models**), we trained Performer models using only the 25%, 50%, 75%, or 100% of the training data, with each larger group inclusive of the previous smaller group. Individuals used for tuning and testing were not downsampled.

Three model replicates were trained for each gene and downsampled group, and the mean of PCC and R^2^ over the replicates was computed. We used 40 genes for this analysis, selecting 20 genes with the highest elastic net R^2^ (averaged over three model replicates) from our initial set of 301 Whole Blood training genes (see, **Selection of train and test genes**) and 20 other random genes from the same set. Performer models were trained using GTEx Whole Blood data.

### Predicting average expression of test genes

We calculated the average Whole Blood TPM, among all 670 GTEx individuals with WGS and Whole Blood RNA-seq data, separately for each of the 1398 genes in the test set. Then, for each of these genes, we passed the 49,152-bp hg38 reference genome TSS-centered sequence and formed a prediction with Enformer, all 903 single-gene Performer models, and all three 300-gene Performer models. Similar to predicting Whole Blood expression from personal genome sequences, we used the ‘CAGE:blood, adult, pool1’ (track 4950) Enformer output and summed over the three TSS-centered bins (191-193).

This left us with one observed and predicted average gene expression value for each gene and model. We took the correlation between the predicted and observed average expression value across the 1398 genes for each model.

### Code and Data Availability

Code to train Performer models and evaluate Performer and Enformer models are available at https://github.com/shirondru/enformer_fine_tuning. GTEx WGS and ROSMAP data are available from dbGaP (accession: phs000424.v9.p2) and Synapse AMP-AD Data Portal (accession: syn2580853), respectively.

## Acknowledgements

We thank the labs of Nilah Ioannidis, Sara Mostafavi, and Olga Troyanskaya for helpful conversations about cross-individual training. This work was funded by an NSF predoctoral fellowship (S.D.) and gifts to K.S.P. from the Biswas Family Foundation, Keck Foundation, Additional Ventures, and the L.K. Whittier Foundation.

