## Supplemental Figures for "Deep-learning prediction of gene expression from personal genomes"

### SUPPLEMENTARY FIGURES

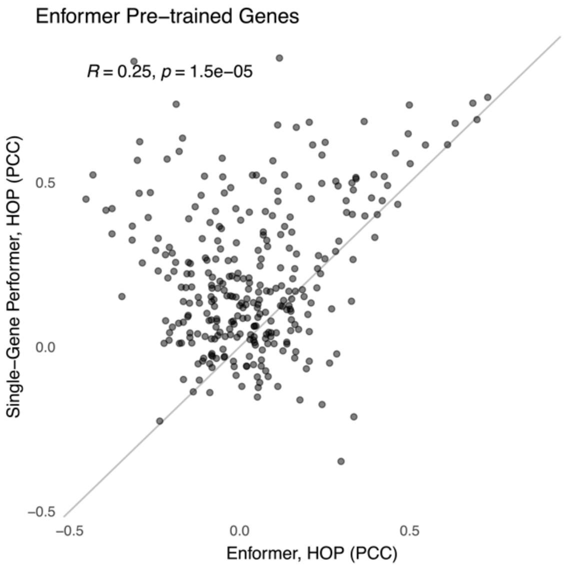


**Figure S1: Fine-tuning improves predictions of expression variability.**
PCC of Performer trained on single genes (y-axis) versus Enformer (x-axis), both evaluated on HOP for each of the 301 training genes in Whole Blood. For most genes, Performer explains more expression variability and is negatively correlated with observed expression less often. $R^2$ and PCC values are averaged as in (**Fig. 1B**).

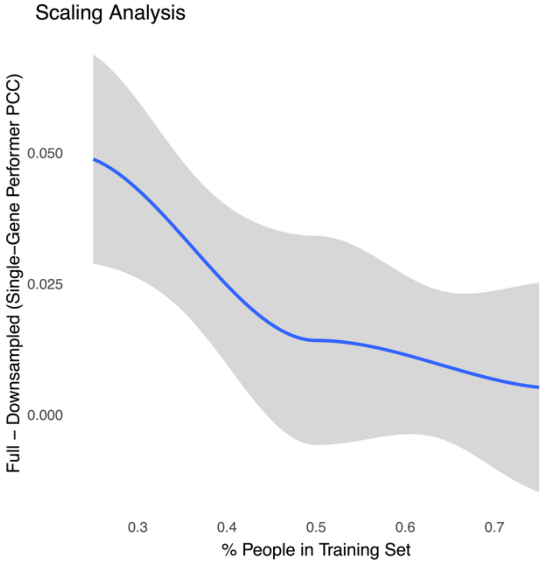


**Figure S2: Fine-tuning performance while downsampling donors used to train.**
Performer models were trained on blood RNA-seq of 40 genes after downsampling the 536 train set individuals (**Methods**). Y-axis represents the decrease in PCC after using 25%, 50%, or 75% of individuals, relative to when training with 100% of the training set. PCC values are averaged over three model replicates for each downsampling group as in (**Fig. 1B**) We observe performance with 50% of the training set (~250 individuals) is comparable to performance after training on the full set, and the trend does not suggest performance would further increase if more individuals were available. Models are evaluated on HOP, using the full set of test individuals. Grey area represents the 95% confidence interval of a LOESS curve fit across genes.

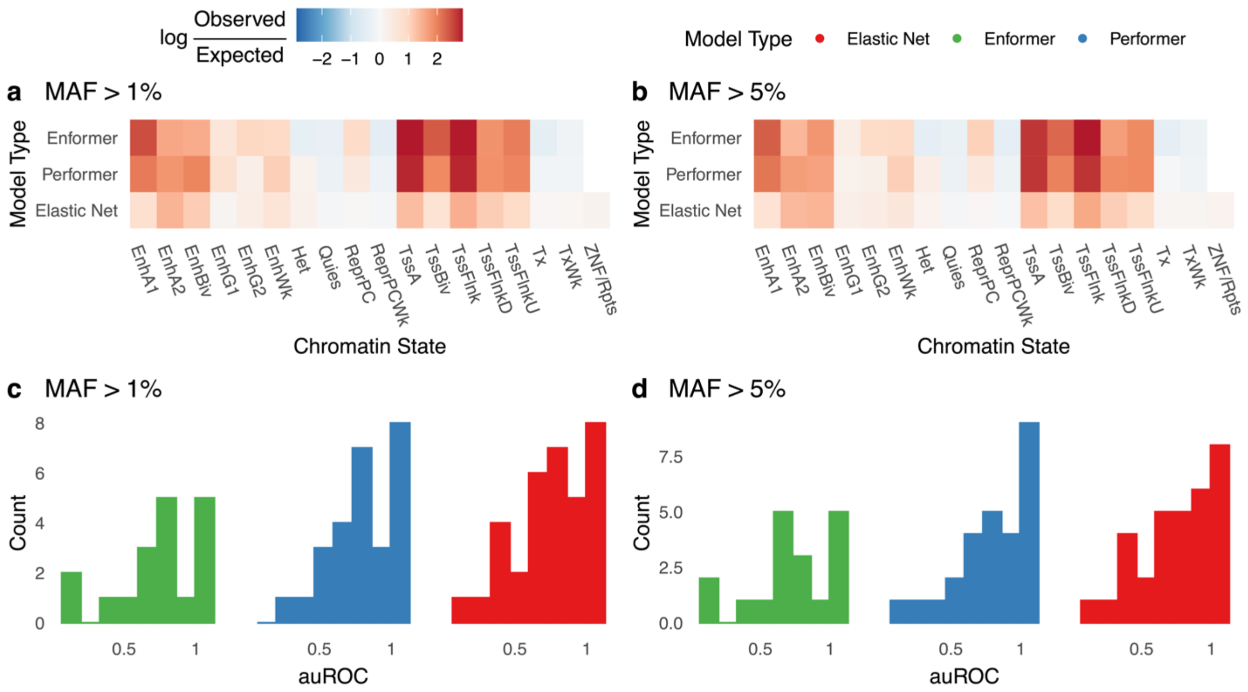


**Figure S3: Functional signatures of HSVs after removing rare variants.**

a) Enrichment of HSVs with minor allele frequency (MAF) >1% across ChromHMM chromatin states (as in **Fig. 3A**).

b) Distribution of area under the Receiver Operating Characteristic (auROC) curve for predicting fine-mapped Whole Blood GTEx eQTLs using HSVs with MAF >1% (as in **Fig. 3C**).

c) Enrichment of HSVs with minor allele frequency (MAF) >5% across ChromHMM chromatin states (as in **Fig. 3A**).

d) Distribution of area under the Receiver Operating Characteristic (auROC) curve for predicting fine-mapped Whole Blood GTEx eQTLs using HSVs with MAF >5% (as in **Fig. 3C**).

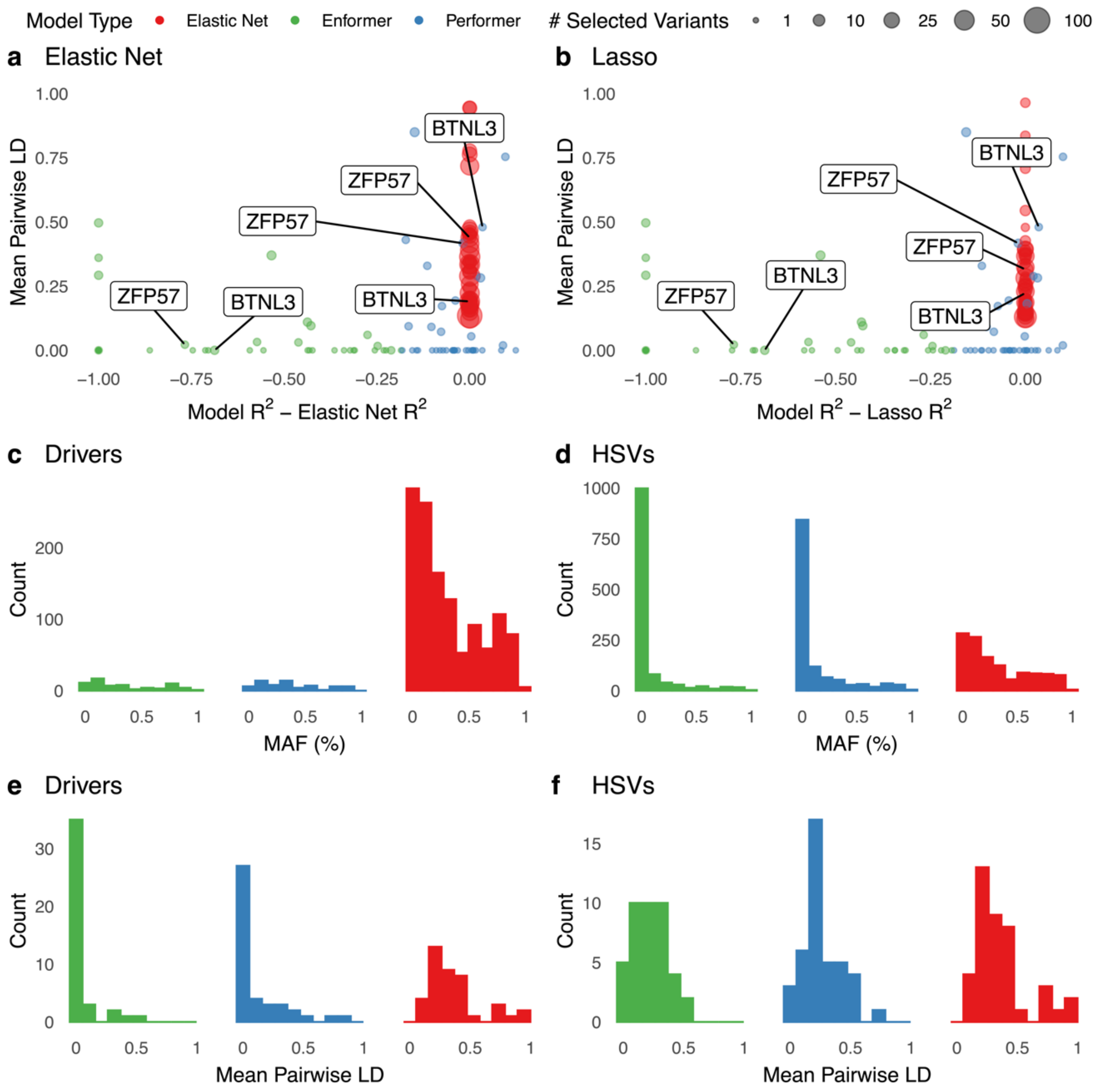


**Figure S4: Properties of driver variants.**

All panels analyzed on 42/301 genes with Performer $R^2 > 0.2$ in Whole Blood**.**

a) Model performance, evaluated on HOP, relative to elastic net plotted against the mean pairwise linkage disequilibrium (LD) of selected variants, with the number of selected variants (Enformer/Performer: drivers (**Methods**), elastic net: non-zero coefficients) indicated by size. Identical to **Fig. 4E**.

b) Same as (**A**) but fitting a lasso linear model.

c) Distribution of minor allele frequency (MAF) for Enformer and Performer driver variants, and elastic net non-zero coefficients.

d) Distribution of minor allele frequency (MAF) for all Enformer and Performer HSVs, and elastic net non-zero coefficients.

e) Mean pairwise LD of Enformer and Performer driver variants and elastic net non-zero coefficients.

f) Mean pairwise LD of all Enformer and Performer HSVs and elastic net non-zero coefficients.

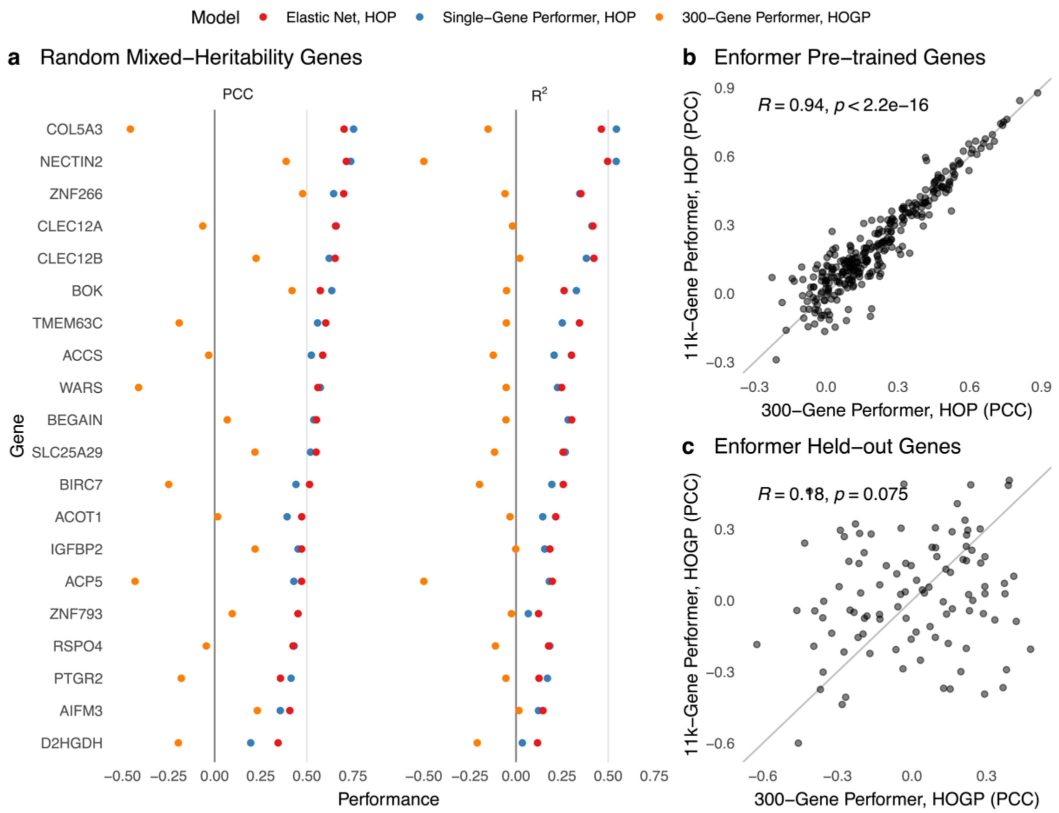


**Figure S5**: **Performance of multi-gene Performer models on Train and Test Genes.**
Each point represents performance of a model evaluated on a single gene using GTEx Whole Blood expression values. Results are not averaged over replicates as in (**Fig. 1B**); they come from one model and one evaluation set only.

a) $R^2$ and PCC of elastic net (red) and single-gene Performer models (blue) that were trained individually on 100 genes in the test set and evaluated on these same genes and HOP. Performer models trained jointly on 301 train genes and then evaluated on HOGP using these 100 test genes (orange) underperform on the 100 unseen genes relative to the models trained directly on those genes. This panel shows 20/100 representative test genes with a range of cis-heritabilities. 300-gene Performer occasionally exhibits strong PCC yet poor $R^2$ on these HOGP because it ranks individuals well but predicts much smaller than observed gene expression changes.
b) PCC of a Performer model trained jointly on 301 genes and evaluated on each of these genes using HOP (x-axis) and of a Performer model trained jointly on 11,429 genes (including the same 301 genes) and evaluated in the same way on the same genes. Each point represents one of the 301 genes used to train both models. Training on more genes did not improve performance on training genes.
c) PCC of the same multi-gene models as in (b) evaluated on HOGP using 100 test genes. Each point represents one test gene that was held-out from training of both models. Training on more genes did not improve performance on unseen genes.

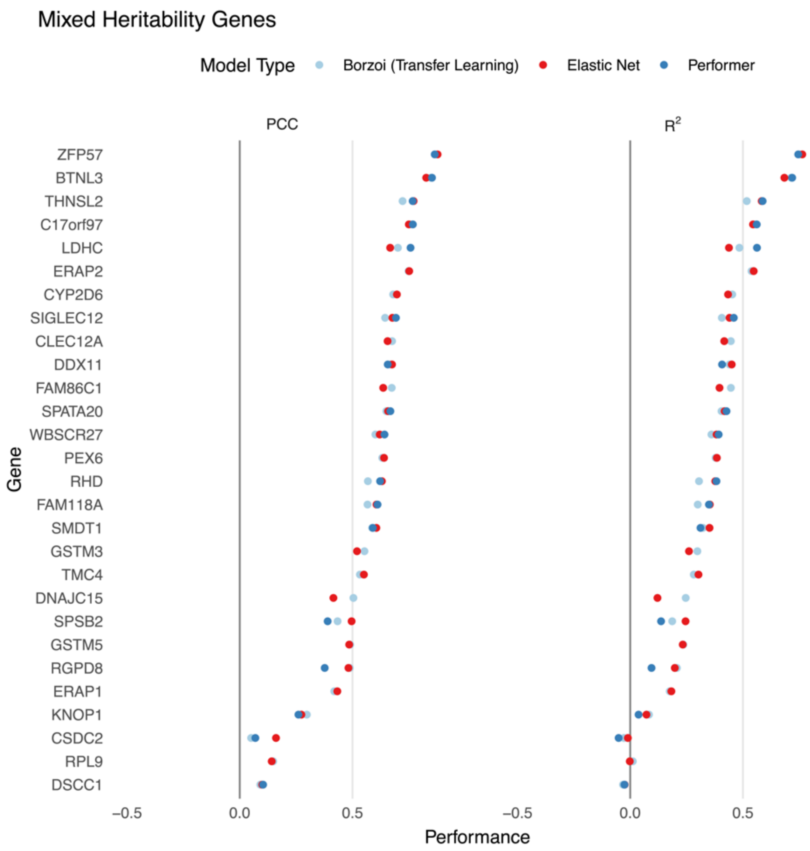


**Figure S6: Performance of Borzoi after transfer learning.**
$R^2$ and PCC of models trained on single genes using GTEx Whole Blood data. All models were trained on single genes and evaluated on HOP. Results come from one model trained on each gene. Elastic net, fine-tuned Enformer (Performer), and Borzoi models after transfer learning (model weights frozen, output layer trained) perform comparably.

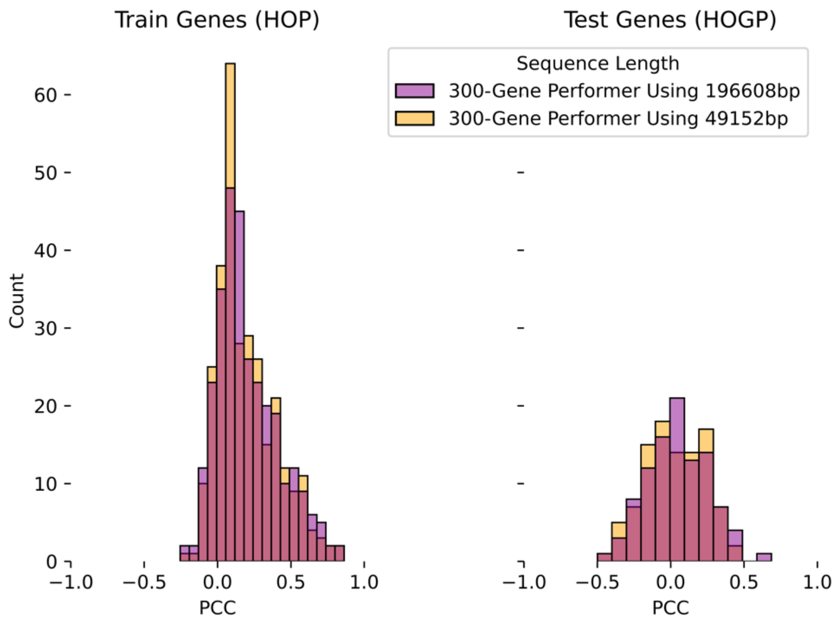


**Figure S7**: **Fine-tuning on longer sequences does not enable better cross-individual performance**. PCC of Performer models trained jointly on 301 genes using GTEx Whole Blood data and either 196-kb DNA sequences (orange) or 49-kb sequences (blue) centered on each gene's TSS, evaluated on HOP using the same 301 train genes (left) or HOGP using 100 test genes (right). PCC values are averaged over three model replicates, as in (**Fig. 1B**).

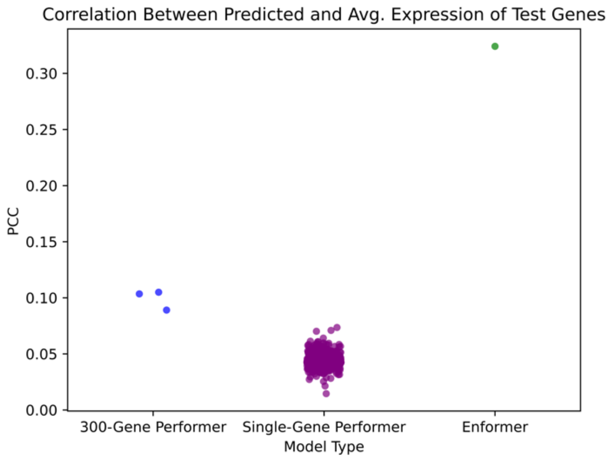


**Figure S8**: **Fine-tuning decreases performance on average expression of unseen genes.**
PCC between average expression and predicted expression, across 1398 Whole Blood genes held out from training of each model. Models included three replicates of a multi-gene Performer model trained on 301 genes (3 models), three replicates of single-gene Performer models trained on each of these 301 genes (903 models), and Enformer. Predictions come from 49-kb reference genome (hg38) sequences centered on each gene's TSS.
